# Early outcomes of England’s new biodiversity offset market

**DOI:** 10.1101/2025.06.22.660961

**Authors:** N.E. Duffus, S.O.S.E. zu Ermgassen, R. Grenyer, O.T. Lewis

## Abstract

Nature markets are increasingly being operationalised as a tool for filling the biodiversity finance gap to achieve global biodiversity targets. One example is the off-site market underpinning the requirement for construction projects to achieve Biodiversity Net Gain (BNG) in England. Here, we quantify the outcomes of the market after 16 months of operation, collecting data from England’s offset register and planning portals. We evaluate the habitat diversity being delivered and how it aligns with environmental targets and existing efforts to achieve goals such as 30×30. Current habitat delivery is skewed toward a few habitat types, with ‘other neutral grassland’ constituting 44% of planned habitat. There is very limited delivery of new wetland and woodland habitats, with most commitments being to improve the condition of existing habitats. Perhaps more importantly, only around 2% of the registered offset area has been sold. Our results demonstrate how the design of the statutory biodiversity metric used to measure BNG is shaping the market through incentives within the metric. The results also give insight into the demand for off-site BNG units in England and the role which government regulation plays in setting demand and therefore scaling the market.

## 1. Introduction

Countries around the world have committed to high-level targets to conserve and restore biodiversity under the Kunming-Montreal Global Biodiversity Framework (KMGBF) (Convention on Biological Diversity, 2023). However, countries face a challenge in financing these activities, with an estimated ∼$700 billion biodiversity finance gap (Deutz et al., 2020). To address this gap, target 19 of the KMGBF aims to mobilise $200 billion per year for biodiversity from all sources, including by leveraging private finance (Convention on Biological Diversity, 2023). Nature markets are important instruments for channelling private finance into biodiversity (Seidl et al., 2024). Nature markets involve creating a commodity from a dimension (or dimensions) of biodiversity, such as habitat area, species richness, or species abundance, and facilitating the sale of this commodity within a market. These markets may be compliance based, allowing buyers to meet a regulatory biodiversity offsetting requirement, or they may be voluntary commitments (Koh et al., 2019; Wunder et al., 2024).

In England, there are ambitious plans to use nature markets to finance the commitments to the KMGBF, with a target to mobilise £1 billion toward achieving England’s Environmental Improvement Plan (EIP) by 2027 (Defra, 2023). Operational nature-based markets in England include the Woodland Carbon Code, the off-site Biodiversity Net Gain (BNG) market, the nutrient neutrality market, the Peatland Code, with new markets proposed for saltmarsh, seagrass, and soil carbon (Defra, 2023).

The off-site BNG market is a new and rapidly scaling market, underpinning the delivery of England’s BNG policy requirement (*Environment Act 2021*). From early 2024, BNG has required new construction projects to deliver a 10% BNG post-development, secured for 30 years, and measured using the statutory biodiversity metric (Defra, 2024b). The statutory biodiversity metric measures the baseline biodiversity value in ‘units’ using a calculation based upon habitat area, distinctiveness, condition, and strategic significance. Distinctiveness is derived from the conservation value of a given habitat type, and condition is derived from a simple checklist of primarily vegetation features (Defra, 2024b). Strategic significance affords more units to habitats which deliver the mapped objectives of a Local Nature Recovery Strategy (LNRS). The biodiversity unit value must be increased 10% post-development through the enhancement and creation of habitats, measured using the same metric but with further multipliers for difficulty, time, and location. More biodiversity units are afforded to habitats which are easy to create, mature rapidly, and are closer to the site of impact. These multipliers are intended to disincentivise developers from damaging habitats which are difficult to replace or take a long time to mature, by requiring them to purchase a larger area of these habitats to compensate for the loss.

BNG is achieved by following the Biodiversity Gain Hierarchy. Preference is given to habitats delivered ‘on-site’ within the development footprint, with any remaining unit liability being secured ‘off-site’ via the off-site BNG market (DHCLG, 2024). As a last resort, where units cannot be found on the off-site market, developers can purchase statutory biodiversity credits from the Secretary of State to meet their BNG liability (DHCLG, 2024). The off-site market allows landowners to enhance and create habitats on their land and then sell the created biodiversity units to developers. To sell units, landowners must enter a legal agreement with either a Local Planning Authority (LPA) or a designated responsible body and have their site listed on the Biodiversity Gain Site Register (Defra, 2024c). The transactions between developments and offsets are also recorded on this public register (Defra, 2025a).

BNG in England has been projected to deliver up to 5428 hectares of habitat for wildlife annually (Defra, 2019) and off-site BNG sites are intended to contribute directly toward the KMGBF target to protect 30% of land by 2030 (Defra, 2024a). Excess habitat creation under BNG can also contribute to the domestic target to create 500,000ha of wildlife-rich habitat by 2042 (Natural England, 2024). Although BNG is a habitat-based approach, it is also expected that the enhancement and creation of habitats will have benefits for species, thereby also contributing to targets to halt and reverse the decline in wildlife populations (Defra, 2023). Therefore, the off-site BNG market is expected to make an important contribution to important domestic and international biodiversity targets.

A potential limitation of using market-based mechanisms for biodiversity conservation and restoration efforts is that, they can lead to the delivery of what is incentivised in the design of the market, rather than what is ecologically desirable (Carver & Sullivan, 2017; Robertson, 2006). Nature-based markets are frequently optimised to deliver what is being measured or commodified. For example, the Woodland Carbon Code measures carbon and therefore, the creation of woodland types which deliver the highest quantity of carbon are incentivised (Stanley, 2024) rather than the woodland type which is most ecologically appropriate for a given location. These incentives have also been observed in the US Wetland Mitigation banking system where project proponents are strongly incentivised to use barrier removals over other restoration measures (Theis & Poesch, 2024). In the case of the off-site BNG market, the habitat-based statutory biodiversity metric may also be creating incentives. In simulations of habitat transitions from common baselines, the multipliers within the metric may incentivise the creation of habitats which are easy to create and mature rapidly (Miles et al., 2025). However, it is not clear how the metric score interacts with the profitability of biodiversity units of different habitat types and the strategic significance multiplier which should incentivise habitat delivery in line with the LNRS objectives. There has not yet been a data-based analysis of what which habitats are potentially being incentivised in practice, or how this delivery complements or aligns with efforts to achieve biodiversity conservation and restoration targets.

Furthermore, the supply-demand dynamics have not yet been explored. In a sample of early-adopter councils, the preference for on-site BNG led to more than 90% of biodiversity units being delivered on-site, with the very small off-site component delivered primarily from purchases by small developments which could not readily achieve their BNG entirely on-site (Rampling et al., 2023). It is not yet known whether this trend is the same now that BNG is mandatory nationally, but it is of substantial interest given the requirement for a reliable source of demand to support the viability of the off-site BNG market and its contribution to financing biodiversity.

Here, we evaluate the outcomes of the off-site BNG market after 16 months of the policy being mandatory. We explore the habitat types being delivered on the off-site market, their potential contribution to environmental targets in England, and how they complement other efforts to achieve 30×30. We also evaluate the demand for off-site BNG units so far, and the implications for future delivery of habitats via the off-site BNG market.

## 2. Methods

### 2.1 Data preparation

#### 2.11 Biodiversity offset data

Information on the occurrence of biodiversity offsets registered between February 2024 and the 31^st^ May 2025 were retrieved from the publicly available Biodiversity Gain Site Register (Defra, 2025a). Kujala et al., (2022) lays out the standard required for offset registers to be sufficiently transparent to track the offset outcomes. The gain site register does not fulfil several of these criteria e.g., by not providing shapefiles of where offsets are located. Therefore, we used the following methods to create our own dataset. However, there are further limitations to the register which cannot be overcome. For example, the register does not match up baseline and post intervention parcels, making it impossible to calculate estimates of the biodiversity units generated by each offset. Furthermore, on the transaction side, where multiple habitat types are purchased, the record for biodiversity units is not parsed out by habitat type (as area is), making it impossible to know how many biodiversity units have been sold per habitat type.

The grid reference provided on the register was used to locate the offset in QGIS and a polygon of the red line boundary was drawn using the map provided by the register as a reference (QGIS Development Team, 2025). The baseline and post intervention habitat data for each offset was converted to a CSV file corresponding to the red line boundary polygon. For each baseline and post-intervention habitat parcel or linear feature we recorded the broad habitat type, specific habitat type, habitat distinctiveness, habitat condition, and area (ha) or length (km). The type of legal agreement (either Section 106 or Conservation Covenant) and the Local Planning Authority (LPA) or responsible body involved in the agreement was also extracted. The Natural England map of National Character Areas was used to obtain data on the National Character Area (NCA) and LPA for each offset (Natural England, 2025b). In total our analyses included 84 offset projects covering 2818.68 hectares, with 1686 baseline parcels and 1593 post-intervention parcels.

#### 2.12 Development transactions data

Information on transactions between developments and offsets was also extracted from the Biodiversity Gain Site Register under the allocation tab for each offset (Defra, 2025a). We extracted the information for every transaction that occurred between February 2024 and May 2025, totalling 234 sales of habitat units. From the off-site register, we recorded the LPA and NCA of the development, offset ID the purchase was made from, habitat type and condition purchased, area (ha) purchased, and the number of biodiversity units purchased. For the transactions between February 2024 and February 2025, we also used the provided LPA and planning reference number to access the planning application on the planning portal for that LPA. The red line boundary of the development was taken from the planning application, digitised into QGIS as a separate shapefile layer and the area of the development was calculated. This gave us a dataset of 76 developments which had purchased off-site units in the first year of the market’s operation.

#### 2.13 Protected area data

We acquired shapefiles for National Nature Reserves (NNRs), Special Areas of Conservation (SACs), Special Protection Area (SPAs), and Sites of Special Scientific Interest (SSSI) from the Natural England Open Data Geoportal (Natural England, 2025a), in line with England’s 30×30 inclusion categories (Defra, 2024a). We excluded National Parks, National Landscapes, Ramsar sites, and The Broads from the analysis owing to their less strict conservation designation than NNRs, SACs, SPAs, and SSSIs.

The shapefiles were merged into a single protected area network layer in QGIS. Since protected area designations frequently overlapped, the ‘dissolve’ and ‘multi-part to single part’ geoprocessing tools in QGIS were used to remove any overlap to avoid double counting of areas. The ‘sf’ package in R (Pebesma & Bivand, 2018; R Core Team, 2024) was used to clip the datasets to the boundaries of low-water terrestrial England, using the boundary from the Open Geography Portal from the Office of National Statistics (Office for National Statistics, 2025). For further analysis, the habitat type of the protected area polygons was classified using the Priority Habitat Inventory (PHI) (Natural England, 2025a) or the Centre for Ecology Land Cover map 2023 (UK CEH, 2023), with the PHI where available taking priority for classification as it provides greater detail for scarcer habitats.

### 2.2 Analysis

#### 2.21 Offset habitat transitions

The area of baseline and post-intervention habitat was grouped by habitat type and condition and plotted.

#### 2.22 Habitat complementarity

The habitat complementarity of the offset network and the protected area network was calculated in R, with habitat complementarity defined as the extent to which the off-site BNG sites supplement the coverage and representation of habitat types within the protected area network. This analysis was conducted to compare the habitat coverage of off-site BNG with the habitat coverage of the protected area network. The complementarity metric for each habitat type was calculated following Palfrey et al., 2022:

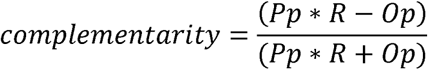

where Pp is equal to the percentage of a given habitat type’s total area conserved within BNG offsets, R is the area of the protected area network divided by the area of BNG offsets, and Op is the percentage of a given habitat type’s total area conserved by protected areas. The complementarity metric generates a value of between −1 and 1, with negative values indicating less than expected habitat complementarity and positive values more than expected. The area of habitat types conserved by offsets and protected areas was extracted from the data prepared above, while the total area of each habitat type in England was extracted from the PHI and CEH land class map, clipped to the England mean low water mark.

#### 2.23 Offset transactions

Descriptive statistics on transactions were generated in R for the volume of unit sales, habitat area sold, the development area, and the relationship between the LPA/NCA of offsets and developments.

## 3. Results

### 3.1 Habitat transitions under BNG

Across the 84 offsets investigated, 37 habitat types were enhanced or created, from 50 baseline habitat types (Supplementary Table 1). The baseline habitat types were predominantly arable, with 34.62% coming from cropland and horticulture and 25.83% from modified grassland, mostly in poor condition (Fig 1a). The planned habitat scenario was dominated (44.22%) by a single habitat type, other neutral grassland, 74.31% of which will be good condition (Fig 1b). Behind other neutral grassland, mixed scrub (13.22%), lowland meadow (11.20%), blanket bog (3.95%), and lowland mixed deciduous woodland (3.87%) are the habitats delivered in the highest quantities.

**Figure 1.**
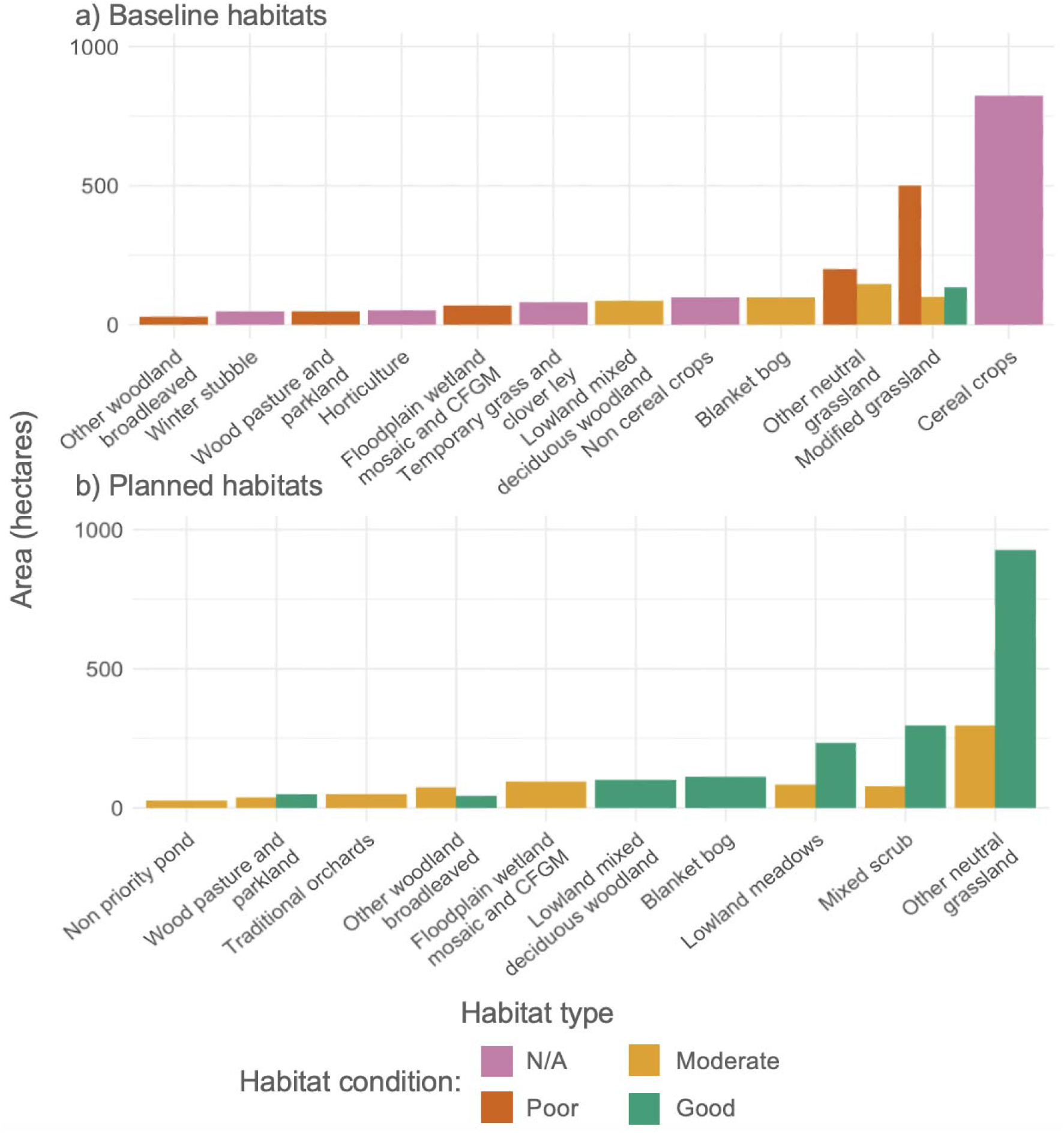
The 15 most common baseline habitats by area (a) and planned habitats by area (b) on the Biodiversity Gain Site Register.

Overall, 64.79% of planned habitat area is grassland, 14.96% is heathland, 13.02% is woodland, and 4.40% is wetland. There are also differences among broad habitat types in the amount of enhancement to existing habitats versus creation of a new habitat e.g., from a cropland baseline. There is a 553.33ha net increase to the total area of grasslands and 378.80ha increase to the area of heathlands, indicating that this quantity of habitat is new (Fig 2). The overall net increase to woodland is 116.14ha and to wetland is just 1.06ha. Many of the commitments to provide woodland and wetland habitats are therefore improvements to the condition of existing habitat.

**Figure 2.**
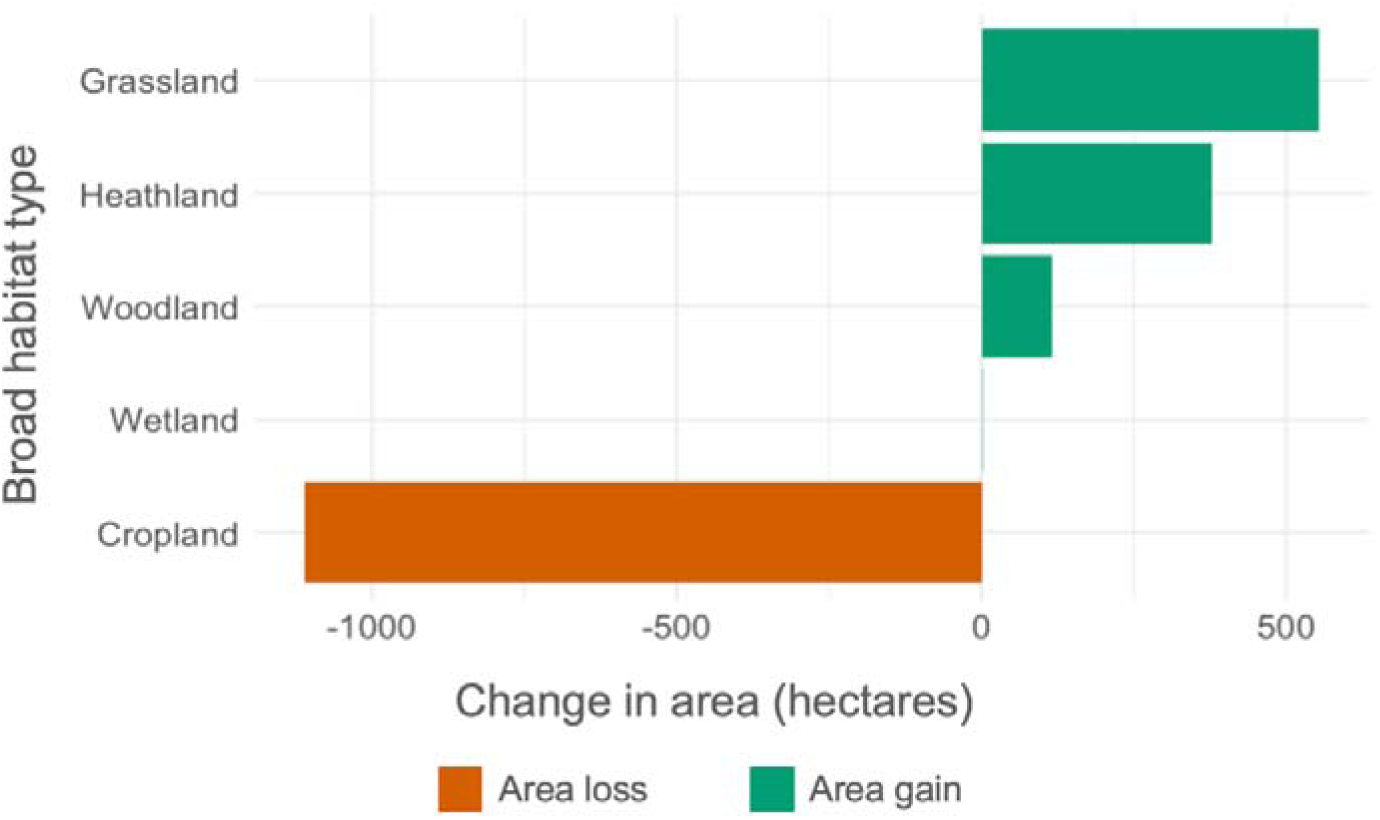
Net changes in area of the five most common broad habitat types on the Biodiversity Gain Site Register.

Although there was substantial creation of new grassland and heathland, these were dominated by other neutral grassland, lowland meadow, and mixed scrub habitat types, with very little creation of other habitats under these broad categories. For example, just 1.11ha of new grassland creation was calcareous grassland and just 40.03ha of new heathland creation was upland or lowland heathland.

### 3.2 Habitat complementarity with protected area network

Offsets were found to have greater than expected complementarity and supplement the habitat coverage of the protected area network for acid grassland (0.11), lowland meadow (0.76), neutral grassland (0.96), traditional orchards (0.70), and upland meadows (0.17) (Fig 3). Complementarity was less than expected for all other habitats and was the lowest for bog (−0.77), calcareous grassland (−0.83), and coniferous woodland (−0.99). This demonstrates that offsets increase the habitat coverage of these grassland habitat types, complementing the existing coverage of the protected area network. However, for all other habitat types, the offset network does not complement the existing coverage of the protected area network.

**Figure 3.**
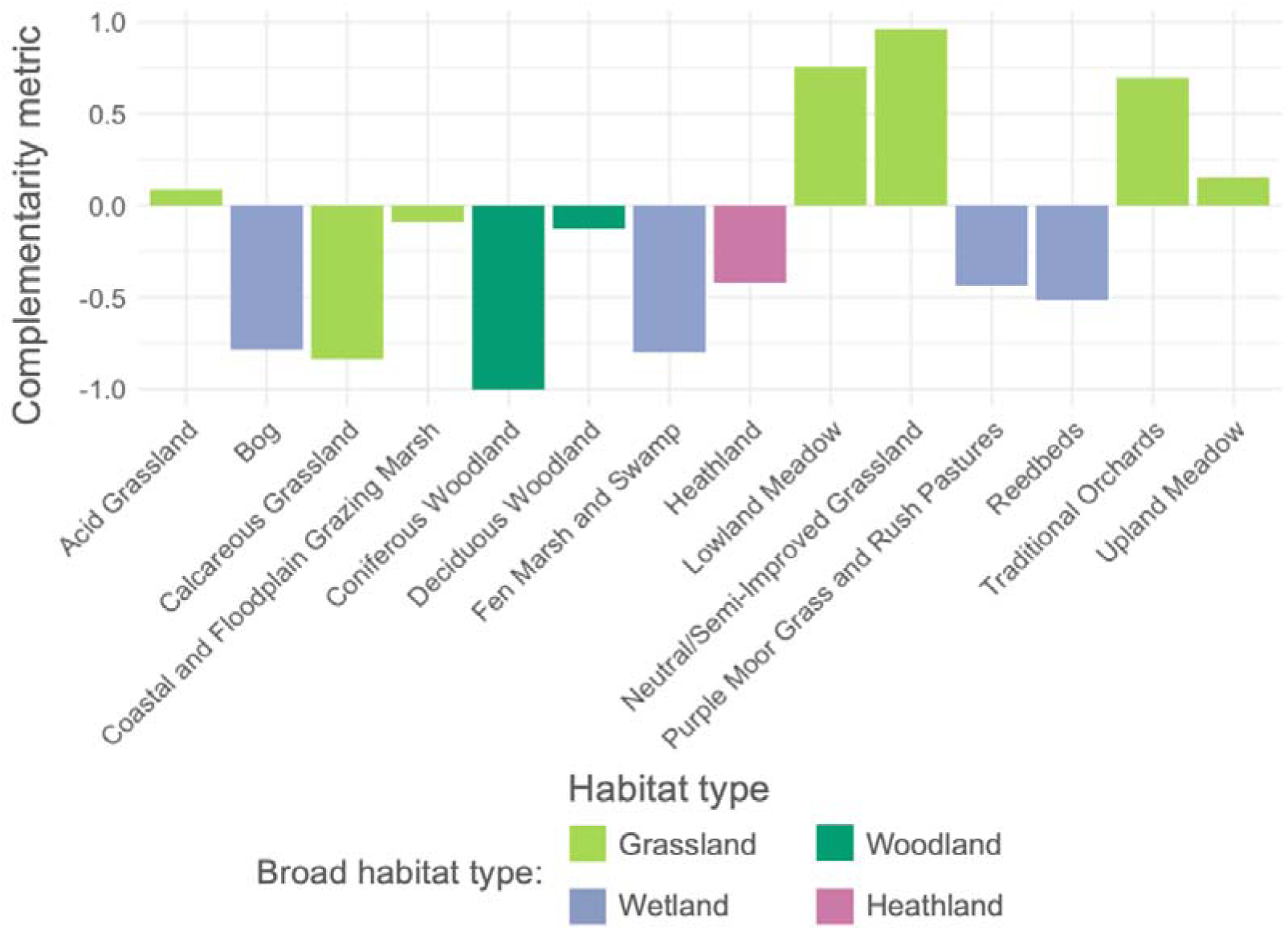
Habitat complementarity metric values for habitat types between off-site BNG and the protected area network in England.

### 3.3 Offset transactions

Between February 2024 and May 2025, a total of 286.77 habitat units were sold to developments across 234 transactions. These units equated to a habitat area of 58.30ha (Supplementary Table 2). This is 2.08% of the land currently registered on the off-site register. The mean transaction was extremely small: 1.22 biodiversity units or 0.19ha of habitat. From the 36 habitat types being offered on the gain site register, only 10 had sold units so far. The most frequently bought habitat was other neutral grassland, constituting 32.30ha of the purchased habitat, followed by lowland meadows (9.08ha), mixed scrub (7.60ha), and floodplain wetland mosaic and CFGM (3.63ha) (Fig 4). For these transactions, we also looked at the spatial penalty incurred. No spatial penalty was incurred for 38.03% of developments, as they purchased from offsets in the same LPA or NCA. The full spatial penalty applied to 35.47% of developments which purchased from offsets from non-adjacent LPA/NCAs, with these developments therefore requiring 50% more units. The remaining 26.41% of developments purchased their units from an adjacent LPA or NCA, therefore requiring 25% more units to meet their BNG liability.

**Figure 4.**
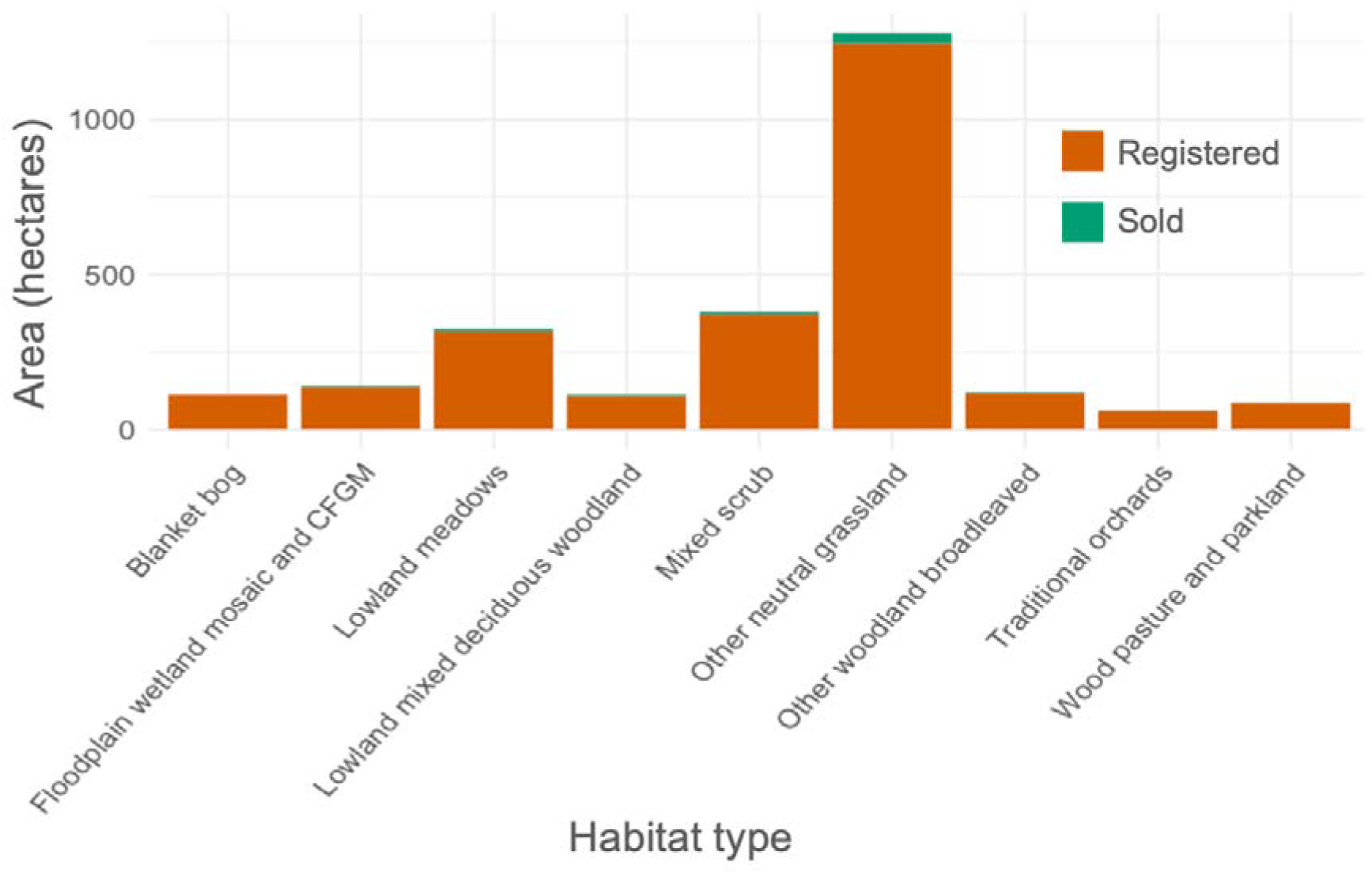
Area of habitat registered on the Gain Site Register and area of that habitat sold to development for 9 common habitat types.

The size of development purchasing units between February 2024 and 2025 was also evaluated. The mean area of developments purchasing off-site BNG units was 0.96ha. Of the developments purchasing off-site units, 70% were less than 1ha in area. However, these smaller sites bought fewer units, with sites less than 1ha purchasing 38.3% of the units sold on the market.

## 4. Discussion

In the first 16 months of mandatory BNG in England there have been substantial commitments to enhance and create habitats to provide biodiversity offsets for developments to purchase. However, our results show that there is currently a substantial skew in the habitats being promised for offsite BNG. Almost half of the planned habitat enhancement and creation is a single habitat type – other neutral grassland. In England, these are already common, widespread grasslands found on neutral soil with a typical floristic diversity of >8 species per m^2^, with several different sub-communities falling under this habitat type (UKHab, 2023). The delivery of this grassland, alongside other grassland types such as lowland meadow, is highly complementary to the protected area network, increasing the coverage of grassland in the network. Good condition other neutral grassland also qualifies as a ‘wildlife-rich habitat’ under the definition for the target to deliver 500,000ha of wildlife-rich habitat (Natural England, 2024). However, the utility of other neutral grassland to wildlife is dependent on factors beyond just the habitat type and condition defined within the statutory biodiversity metric (Duffus et al., 2024).

Whilst well-managed other neutral grassland can be a valuable habitat for wildlife, this skew towards a single habitat type within the off-site market seems unlikely to lead to optimal ecological outcomes. For example, the commitments to create new woodlands are limited, which means that the off-site market is making a limited contribution to woodland creation targets (Defra, 2023). Furthermore, when we consider England’s species abundance targets, many of the indicator species are woodland specialists such as the lesser spotted woodpecker *Dryobates minor* or the white admiral *Limenitis camilla* (The Environmental Targets Regulations, 2023). However, given that most woodlands in the UK are in poor ecological condition (Woodland Trust, 2025), off-site BNG could be a useful tool for financing improved woodland management.

The high volume of other neutral grassland delivery relative to other habitats could result from a variety of reasons, including the incentives created by the statutory biodiversity metric. From a cropland baseline, other neutral grassland delivers the highest possible number of biodiversity units per hectare, whilst some priority habitats deliver very limited biodiversity units, and even a net loss in units relative to the baseline (Miles et al., 2025). This is due to the difficulty and temporal multipliers within the metric which incentivise the avoidance of impacts on those rare habitats which will be difficult to replace on short timescales. However, this design also has the effect of incentivising the creation of easily created and rapidly maturing habitats. We also observe these potential incentives on-site, where 23% of promised habitat units in early adopting councils were from moderate condition other neutral grassland (Rampling et al., 2023)

Our results also show that other neutral grassland has been the habitat purchased most on the off-site market, demonstrating demand for this habitat type. Low demand for the high and very high distinctiveness habitats suggests that impacts are likely being avoided on these habitats, as like-for-like off-site units are typically required. This does not mean that other neutral grassland is being lost to development, however; the metric has flexible trading rules for low and medium distinctiveness habitats, which mean that other neutral grassland can be used to offset the loss of any low distinctiveness habitat or any medium distinctiveness grassland habitat (Defra, 2024b). The demand for other neutral grassland on the off-site market is likely to compensate for the loss of low distinctiveness habitats such as cropland and modified grassland which are commonly developed upon (Rampling et al., 2023). These flexible trading rules mean that the loss of these habitats could also be compensated for with medium distinctiveness woodlands, wetlands, or heathlands. However, given the abundance of other neutral grassland units and the ease of creation relative to other habitat types, these units can be cheaper than woodland and wetland units for developers (Biodiversity Units UK, 2025).

In addition to the skew in planned habitats, the distribution of baseline habitats is also of interest. There are growing concerns that conservation interventions may be displacing biodiversity impacts and therefore incurring leakage (Balmford et al., 2025). Our results indicate that leakage may need to be considered when evaluating the biodiversity benefits from the off-site BNG market. The baseline habitats are primarily arable and poor condition modified grassland, with ‘modified’ grassland frequently associated with intensive grazing (UKHab, 2023). This means that the off-site BNG market is possibly associated with a loss in cereal crop yield and a reduction in livestock grazing. The extent of the yield losses will depend on factors such as the productivity of the land. Lost yields can incur a substantial biodiversity footprint depending on where they are compensated for (Ball et al., 2024). A reduction in livestock grazing may not necessarily incur leakage, if coupled with the dietary shifts which are in line with the UK’s carbon budget (Climate Change Committee, 2025). It is also probable that many of the habitat banks delivering grasslands will be grazed in the post-intervention scenario for several reasons. Firstly, grazing will allow landowners to maintain the agricultural status of their sites, which has tax benefits (HM Revenue and Customs, 2024). Secondly, an appropriate grazing regime can be very beneficial to groups such as invertebrates, by maintaining habitat heterogeneity (Lyons et al., 2018). This and other potential impacts of leakage arising from the changes in agricultural production across the off-site network are complex and not yet understood.

Although there has been a substantial volume of habitat enhancement and creation legally secured within the first 16 months of BNG, there has been very limited demand. Over this period, only 2% of the habitat area from the gain site register was sold. This result could be an early indication that the demand for off-site BNG units is lower than expected. Importantly, 38.8% of the off-site unit demand came from small developments of less than 1ha. This is an important result, given that there are current proposals to exempt more developments in this category from BNG entirely and to adjust the metric to make it simpler for the remaining developments in this category to achieve BNG entirely on-site (Defra, 2025c). These changes have the potential to further reduce the demand for units from the off-site market. This is of concern because, unless the units are sold, the habitats planned may not be delivered, depending upon the terms of the legal agreement involved. Therefore, the habitat gains from the off-site market cannot be assured until they have been sold to a development. However, the application of BNG to Nationally Significant Infrastructure Projects (NSIPs) from May 2026 has the potential to substantially increase the demand for off-site BNG units (Defra, 2025b).

The early outcomes of England’s off-site BNG market demonstrate important lessons for the scaling of nature markets elsewhere to meet the targets under the KMGBF (Convention on Biological Diversity, 2023). The skew toward the delivery of a few habitat types demonstrates how the design of a nature market and its measurement tool can incentivise the market outcomes, potentially leading to sub-optimal ecological outcomes (Cusworth & Stanley, 2025). This outcome also demonstrates that private finance through market-based mechanisms alone cannot be relied upon to deliver biodiversity targets, with public finance still required, particularly to deliver habitats which take a longer timescale to mature and may be unprofitable through markets (Kedward et al., 2023; Löfqvist et al., 2023). These more risky and less profitable habitat types are potentially better aligned with public finance.

Finally, the case study of off-site BNG in England demonstrates that the scaling of nature markets to is dependent on government policies and rules which set the demand drivers (zu Ermgassen et al., 2025). While markets may be seen as an alternative to politically unfavourable policies which prevent biodiversity loss, the case in England demonstrates that scaling up markets faces the same political constraints as regulation.

## Conclusion

The off-site BNG market in England is a new mechanism for driving investment into nature. Currently, the market is heavily skewed toward the delivery of easy to create and rapidly maturing grasslands, with limited delivery of more complex or longer maturing habitats. This is probably due to incentives created by the design of the statutory biodiversity metric and the profitability of different habitat types in the market. These incentives limit the potential contribution of the off-site BNG market toward key biodiversity targets. Another potential limitation to the upscaling of the off-site BNG market toward key biodiversity targets are the potential constraints on demand for off-site units created by government intervention in the market. The early outcomes observed from the off-site BNG market have important implications for the scaling of similar nature markets elsewhere to meet the KMGBF.

